# Harnessing Big Data for Systems Pharmacology

**DOI:** 10.1101/077115

**Authors:** Lei Xie, Eli J. Draizen, Philip E. Bourne

## Abstract

Systems pharmacology aims to holistically understand genetic, molecular, cellular, organismal, and environmental mechanisms of drug actions through developing mechanistic or predictive models. Data-driven modeling plays a central role in systems pharmacology, and has already enabled biologists to generate novel hypotheses. However, more is needed. The drug response is associated with genetic/epigenetic variants and environmental factors, is coupled with molecular conformational dynamics, is affected by possible off-targets, is modulated by the complex interplay of biological networks, and is dependent on pharmacokinetics. Thus, in order to gain a comprehensive understanding of drug actions, systems pharmacology requires integration of models across data modalities, methodologies, organismal hierarchies, and species. This imposes a great challenge on model management, integration, and translation. Here, we discuss several upcoming issues in systems pharmacology and potential solutions to them using big data technology. It will allow systems pharmacology modeling to be findable, accessible, interoperable, reusable, reliable, interpretable, and actionable.

## 1. Introduction

Drug action is a complex process. A chemical, which can be synthesized, natural, or endogenous, starts its effect on biological systems with its interactions with biomolecules (proteins, DNAs, or RNAs), that is, its targets. In the case of most commonly investigated protein targets, there are many types of interactions that are determined by protein conformational dynamics, from antagonism to partial agonism and inverse agonism, from biased signaling to allosteric modulation (1; 2). The change of functional state of the biomolecule, which results from the binding/unbinding kinetics and thermodynamics, will ultimately drive biological outputs. Moreover, a chemical rarely binds to a single target. Multiple target binding, i.e., polypharmacology, is a common phenomenon (3). Even multiple weak drug-target interactions can collectively have strong effect on the physiological response of an organism (4). To understand how the modification of the functional state of multiple biomolecules leads to the alteration of the cellular state through gene regulation, signaling transduction, and metabolism, and ultimately causes the change of the physiological or pathological state of the individual, systems biology is an essential component to model, simulate, and predict the phenotypic response of drug action (5). The fate of the drug molecule itself is also dependent on the cellular states of transporters and metabolizing enzymes and physiological environments. Individual genetic/epigenetic variations and life histories add great diversity to the drug action. They may not only impose an effect on the thermodynamics and kinetics of drug binding as well as pharmacokinetics, but also rewire the biological network, thereby resulting in a dramatically different drug response. It has been recognized that a reductionist view of drug action is too simple to explain and predict the drug phenotypic response to complex diseases. A holistic understanding of drug action under diverse genetic/epigenetic and environmental factors, from conformational dynamics of drug-target interaction to emergent properties of biological systems, from enzyme reactions to physiological-based pharmacokinetics, is needed (6). Embracing the concept of multi-scale systematic modeling of drug actions by integrating multiple omics data and biological mechanisms prompts the emergence of a new discipline of systems pharmacology (6; 7). The fundamental theme of systems pharmacology is 1) to develop actionable and interpretable mechanistic or predictive models by integrating biological and clinical data at multiple temporal and spatial scales, and 2) to use the output from the model for generating novel hypotheses, discovering new biomedical knowledge, and supporting decision making in drug discovery and clinical practice. Systems pharmacology provides a promising avenue to gain a comprehensive and holistic picture of drug actions under the complex interplay of genetic, molecular, cellular, organismal, and environmental components. Such understanding may fill the current innovation gap in drug discovery (8).

Data-driven modeling plays a central role in systems pharmacology. Recent efforts in high throughput experiments have generated a huge amount of data across the multiple biological scales of the organism, across a wide range of time scales, and across multiple species. These data sets provide unprecedented opportunities for systems pharmacology, but impose great challenges in data processing, management, sharing, and integration (9; 10). The rapid advances in cloud computing, big data technology, and data science have started to clear the hurdles in data-driven modeling for systems pharmacology. The planned US National Strategic Computing Initiative (NSCI) will maximize the benefits of High Performance Computing (HPC). The development of the National Institutes of Health (NIH) data science commons will make biological data findable, accessible, interoperable, and reusable (11). These efforts will significantly enhance the availability and quality of biological data, thereby enhancing the capability of systems pharmacology modeling. Indeed, systems pharmacology models based on the integration of genome-wide, heterogeneous, and dynamic data sets have already shown promises in drug repurposing (12–14), predicting drug side effects (15–18), and developing combination therapy (19) and precision medicine (20).

With the explosion of mathematical and computational models for genomics, molecular dynamics, biological networks, whole cells, tissues, organisms, and populations, we face new challenges in systems pharmacology. How can we efficiently and effectively manage these diverse models, including sharing, reuse, validation, reproducibility, access, and searching? How can we integrate diversified models that are from different resources, based on different methodologies, and at different temporal and spatial scales into a unified, potentially more powerful mechanistic or predictive model that captures the whole spectrum of drug actions? How can we translate mathematical languages or decipher black boxes representing these models into causal-effect relationships or simple rules that can be comprehended by biologists and clinicians, and integrate them with existing knowledge for automated reasoning and decision making? Addressing these challenges will no doubt facilitate harnessing big data for systems pharmacology, and realize the full power of systems pharmacology in drug discovery and clinical practice. In this paper, we will discuss the unsolved issues in the management, integration, and translation of systems pharmacology models, and propose possible solutions to them using big data technology and analytics. It is expected that the paralleled development of high-throughput techniques, cloud computing, data science, and the semantic web will allow systems pharmacology models to be findable, accessible, interoperable, reusable, reliable, interpretable, and actionable.

## 2. Model management

A comprehensive and holistic understanding of drug action requires the integration of diverse models from different data modalities (e.g., single nucleotide variants, copy number variations, methylations, proteomics, transcriptomics, metabolomics, etc.), across multi-scales of cellular organization, from the atomic details of drug-target binding thermodynamics and kinetics to proteome scale drug-target interactions, from functional impact of mutations to emergent properties of biological networks, from cytochrome P450 enzyme reaction to physiologically-based pharmacokinetics. They can be biophysics-based molecular models, machine learning models of molecular interactions, mathematical models of systems biology or pharmacokinetics, or connectome models of the human brain. Even in the same type of model, models can be significantly different. For example, a molecule dynamics (MD) model of protein structure could be a Cα-represented coarse-grained elastic network model or an all-atomic conformational ensemble from microsecond MD simulations. A drug-target interaction model could be a graphic representation that abstracts each protein and drug as a single node, and the interaction between them as an edge (21), whereas, a drugome model includes 3-dimensional structures of drug-target complexes (22). A systems biology model could be represented as a stoichiometric matrix and flux bounds (23), based on ordinary differential equations, or encoded as three-dimensional geometries and partial differential equations (24). Machine learning models could be inferred using different features and base learners such as Deep Neural Network, Support Vector Machine, or Random Forest. Furthermore, the models from different domains are often interleaved. For example, a drug-target binding/unbinding kinetics model could be a combination of elastic net model and machine learning model (25). The diversity of models makes it a non-trivial task for scientists to discover, access, and reuse a model that is beyond their domain of expertise, as well as to integrate multiple models. Moreover, a model alone may not be sufficient for a real-world application; the model is often dependent on multiple data sets. Big data integration is an important topic in systems pharmacology, which has been covered elsewhere (9; 10). Beyond the data challenge, models are strongly coupled with algorithms underlying the model, software that implements the algorithm to execute the model, and tools that process inputs and outputs. Software is often developed in different languages, complied in different operating systems, and changed over time, making its interoperability, reuse, and reproducibility difficult (26). Moreover, software is rarely professional grade that is robust and easy to use, being developed as part of a research, rather than a development process. Innovative model management strategies for systems pharmacology, including but not limited to model storage, transfer, sharing, standardization, and validation, are in dire need.

### Model storage, transfer, and sharing

With the exponential increase of biological data and computing power, the number of systems pharmacology models increases at an even faster pace. For example, understanding of drug binding thermodynamics and kinetics requires detailed study of the conformational dynamics of protein structures. Now it is possible to sample conformational space of a protein structure in the microsecond time-scale and longer using MD simulations (27). MD generates millions of conformations for further analysis − ten times more than all the structures deposited in the Protein Data Bank (PDB) (28) to date.

Heterogeneous network models of chemicals and proteins are very useful to predict genome-scale drug-target interactions. A network model of all chemicals in ChEMBL (29) may have millions of nodes and tens of millions of edges. Similarly, a network models for all sequences in UniProt (30) can quickly reach 100 million nodes. The number of edges will increase exponentially with the increase of nodes. The volume of systems pharmacology models will impose a hurdle for model storage and transfer, making model sharing difficult. By taking advantage of the underlying properties of biological systems (e.g., redundancy), systems pharmacology models can be compressed without loss of information (31–34). For example, a genome-scale metabolic network can be reduced to only essential and synthetically lethal reactions. Furthermore, big data storage and transfer technology, which has already gained substantial attentions in genomics (35), can be applied to large-scale computational models in systems pharmacology.

### Model standardization

In order to make systems pharmacology modeling findable, accessible, interoperable, and reproducible, a prerequisite is to have computational models adhere to a common standard of representation and annotation, including a description of the execution and outcomes of the simulations. A great deal of efforts has been devoted to standardization in systems biology to facilitate collaboration (36). Now nearly all modeling in systems biology follows the suggestion of the MIBBI project (Minimal Information for Biological and Biomedical Investigations) (37). The minimum information required in the annotation of models (MIRIAM) provides guidelines to curate models (38). The MIRIAM registry proposes a connection between ontologies, model format, databases, and tools (39; 40). Ontologies have been developed to describe model structures and components, mathematical formulizations, and simulation algorithms (36). A number of modeling formats have been proposed to encode systems biology models (36). Particularly, the Pharmacometric Mark-up Language PharmML for the representation and exchange of pharmacometric models is under development (41).

The efforts made by the systems biology community should be extended to systems pharmacology, which is even more diversified than systems biology. It requires the development of new modeling ontologies, utilities, and visualizations in a specific domain, as well as protocols and tools that enable communications across domains. For example, to predict how drug inhibition of gene A and a mutation in gene B collectively affect blood pressure, we may need three models: a biophysical model to determine the strength of competitive inhibition of the drug on its target gene A, a machine learning model to predict the functional impact of mutations on gene B (e.g., neutral or deleterious), and a genome-scale metabolic model or a kinetic model that takes the outputs from drug binding and mutation models as inputs. The output of biophysical model is usually in the form of a binding free energy. The output of machine learning model of mutation could be the probability that the mutation is predicted to be deleterious. The input required for the stoichiometric model and kinetic model is different: the stoichiometric model may need to constrain the flux corresponding to the reaction catalyzed by the target of drug or harbored mutation, whereas the kinetic model may need to modify the kinetic parameters of the corresponding reaction. The integration of popular bioinformatics workflow systems such as Galaxy (42) with cloud computing (43) could be a powerful approach to linking diverse models together. However, existing workflow systems do not have sufficient semantic supports to represent and reproduce the communications between models. An ontology is needed to represent common molecular components and their interactions, which may have different representations in different models. For example, a gene can be represented by a structure of the encoded protein in the biophysical model, but by a fragment of DNA sequence in the mutation model. Eventually they need to map to a variable in a mathematical model. Moreover, detailed descriptions of the experimental procedure (e.g., how to convert the binding free energy to a flux constraint) should be encoded in a way that makes it understandable to both machine and humans. It is not possible for such efforts to be successful without an ecosystem that encourages collaborations. The NIH data science commons is addressing this challenge (11).

### Model validation

A more serious problem in the application of systems pharmacology modeling is how to validate models, assess their accuracy and reliability on a new prediction, and define scope of their application domains. Recent debates on the origins of inconsistency of constraint-based network modeling highlights the difficulties in model validation (31; 44; 45). Constraint-based network modeling is a powerful tool in systems pharmacology. It has been applied to predict drug side effect profiles resulting from off-target binding (46), elucidate mechanisms of antibiotics (47), predict personalized drug response (20), and identify drug targets and biomarkers (48). However, the application and reproducibility of constraint-based modeling are hampered by the lack of standard formats for network models and associated tools to parse the models (31; 44; 45). A more fundamental issue is whether an exact arithmetic solver is required to achieve consistent results (31; 44; 45). The standardization efforts in systems biology discussed previously may facilitate addressing above problems. The minimum information about a simulation experiment (MIASE) has been proposed to unambiguously define how to reproduce simulation results (49). Such information should be included in all types of models in systems pharmacology.

Machine-learning based big data analytics is playing an increasingly important roles in systems pharmacology (10). Due to the nature of pharmacological data that are often biased, incomplete, and heterogeneous, plus our limited knowledge of biological systems and human physiology, the machine leaning models generalized using a specific algorithm and one particular set of data may not be applicable to a new case. Thus, the rigorous on-the-fly validation of machine learning models is critical, particularly, in a risk-sensitive domain (e.g., to determine if a cancer patient is sensitive to an experimental anti-cancer drug, or if a lead compound should move into clinical trials, where failure is costly. In this regard, the validation of a machine learning model for systems pharmacology using conventional techniques such as cross-validations or a limited number of wet-lab experiments is not sufficient, as they do not cover a whole pharmaceutical and physiological space. To clearly define the applicable domain of a model, so that a non-expert can have sufficient information to use the model wisely, new standards are needed during the future development of machine learning models for systems pharmacology. Firstly, the model should be constantly evaluated whenever new data become available. It requires that models are associated with computational tools that extract and format new data, as well as metadata that describe the data on validation. Whenever feasible, the model can be retrained in the framework of active learning (50). Secondly, multiple evaluation metrics are needed, as different applications should be measured differently. For example, a ranking is sufficient for a web search, but not for determining if a drug is effective on a patient. Finally, for each prediction, the reliability of the prediction should be estimated, as a new case may fall outside the generalized hypothesis space of a model that is based on biased and incomplete data. A Case-Based Reasoning (CBR) framework may be useful to address this problem, as detailed in the following section.

### Model access and reusability

Even if systems pharmacology models and their associated data, algorithms, and software are well-defined and validated, it may not be easy for a user to find the most suitable models unless all relevant documents and literatures are studied. Several databases have been developed to host computational models relevant to systems pharmacology such as drug-target interaction (51), constraint-based metabolic network models (52), and pharmacometrics models (53). However, several challenges still remain, which hinder the accessibility and usability of systems pharmacology modeling. Firstly, data, software, and models are scattered around the internet. Diverse systems pharmacology models have not been registered in a central place so that they can be easily accessed. Secondly, data, algorithms, and software that are used to build the model are often separated from each other. As discussed previously, data and software that are used to develop systems pharmacology models are inseparable components of those models. Finally, there is no easy way to search for suitable models to use in a systems pharmacology modeling project.

The National Institute of Health (NIH) data science commons (https://datascience.nih.gov/commons) is building a shared virtual space that allows scientists to find, manage, share, use and reuse digital objects (data, software, metadata and workflows). It is a complex ecosystem including a computing environment (e.g., cloud) that supports access, utilization and storage of digital objects, publically available data sets that adhere to commons digital object compliance FAIR (findable, accessible, interoperable and reproducible) principles, software services and tools that enable scalable provisioning of compute resources, interoperability between digital objects within the Commons, discoverability of digital objects, sharing of digital objects between individuals or groups, access to and deployment of scientific analysis tools and pipeline workflows, connectivity with other repositories, registries and resources.

The concept of a common can include systems pharmacology models, each of which is a collection of model representation, data, software packages, and metadata that describe them. As shown in Figure 1, model, data, software, and metadata can be stored in different computers in a cloud computing environment. To make software reusable, big data technology such as Docker (https://www.docker.com/) can be used to wrap the software. Metadata are needed to standardize and characterize components of data, model, and software, define their interactions, and register the model system in the commons.

**Figure 1.**
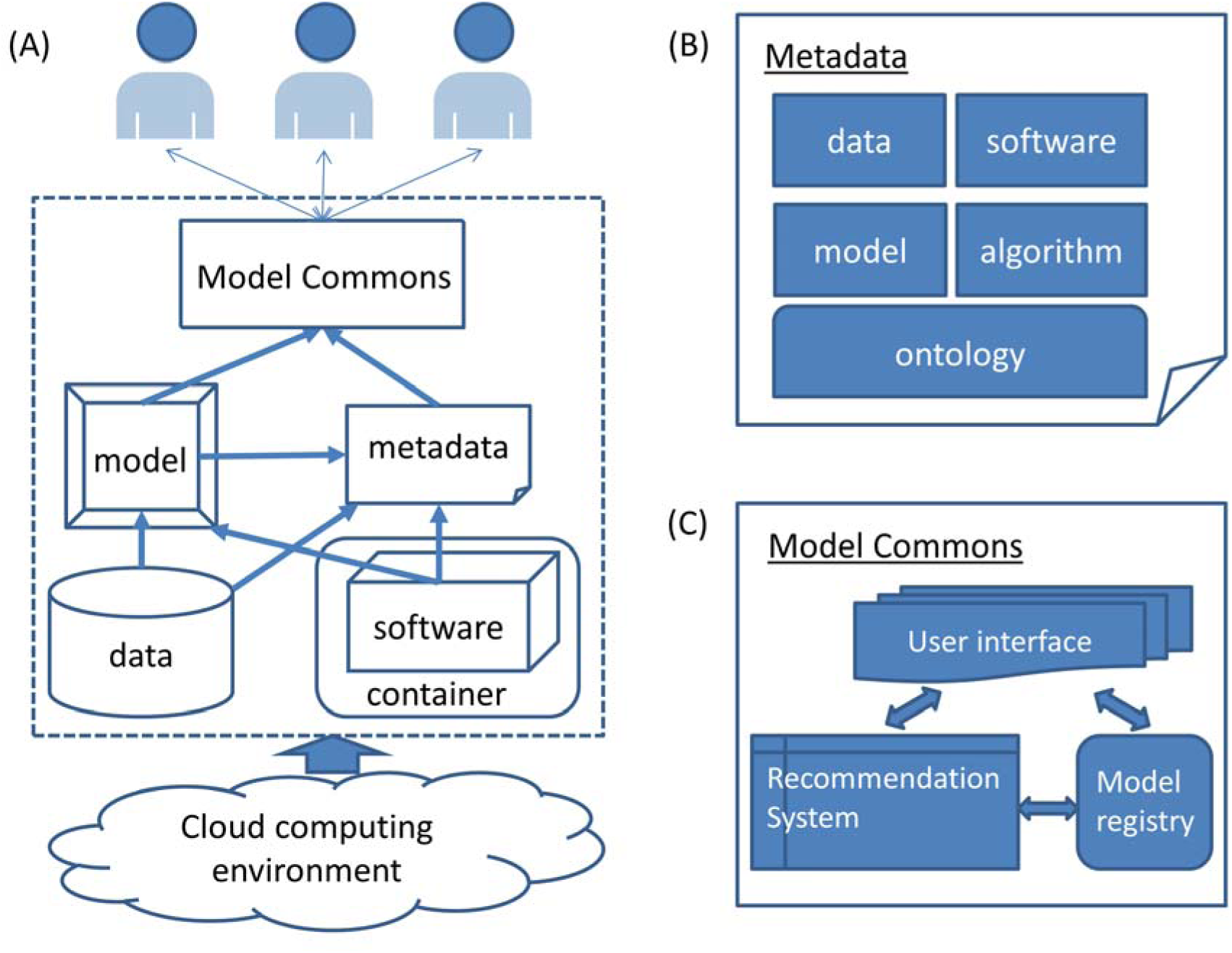
Scheme of a systems pharmacology model management system that adhere to commons digital object compliance FAIR (findable, accessible, interoperable and reproducible) principles. (A) Architecture of model management systems. Models are linked with associated data sets, metadata and software that is wrapped within a container and accessed through a model commons. The whole system may be supported by a cloud computing environment. (B) Model metadata is built on ontologies, including information on the model itself, data, algorithms, and software. (C) The model commons may need a recommendation system to rank the relevant models based on user requests in addition to a model registry and user interfaces.

To make models findable and accessible, a recommendation system similar to those used in Netflix or Amazon can be valuable. Each model should be puts in a global context, such as the usage of model itself, embedded data, underlying algorithm, software package etc in other systems, and applications cited in the literatures. Existing techniques in big data analytics such as pagerank and matrix factorization can be applied to build the recommendation systems using the usage information and the metadata associated with the model. Thus the models can be ranked based on user’s interest.

## 3. Model integration

The integration of multiple diverse models is critical for the success of systems pharmacology modeling. Firstly, it has been recognized that multi-scale modeling is needed for understanding drug actions comprehensively and holistically as well as developing precision medicine (6). The success of multi-scale modeling depends on the mechanistic integration of molecular, network, tissue, organism, and populations models at several temporal and time scales. Due to its complexity, a semantic integration may be needed, as discussed in the next section. Secondly, on the same scale, multiple models can be combined by data fusion, voting, weighting, or averaging. It has been appreciated that the combination of multiple models will generally outperform a single model. However, there is no one-size-fit-all solution to achieve the optimal model combination in the context of systems pharmacology. Analog to the data integration in systems pharmacology (7), we refer to the model integration across different scales as vertical model integration, but model combination at the same scale as horizontal model integration. Finally, one of the fundamental questions in systems pharmacology is how to link *in vitro* drug potency to *in vivo* drug activity, and how to extrapolate the drug response in animal models to that in humans. Solving these problems requires effective methods to reduce the transportability bias. It rises when the population from which data are acquired is different from the one for which the inference is intended. Thus, the success of model integration in systems pharmacology depends on solutions to multi-scale modeling (vertical model integration), horizontal model integration, and model transportability, as shown in Figure 2. The problem of multi-scale modeling and model transportability in systems pharmacology has been discussed elsewhere (10). Here we will focus on horizontal model integration.

**Figure 2.**
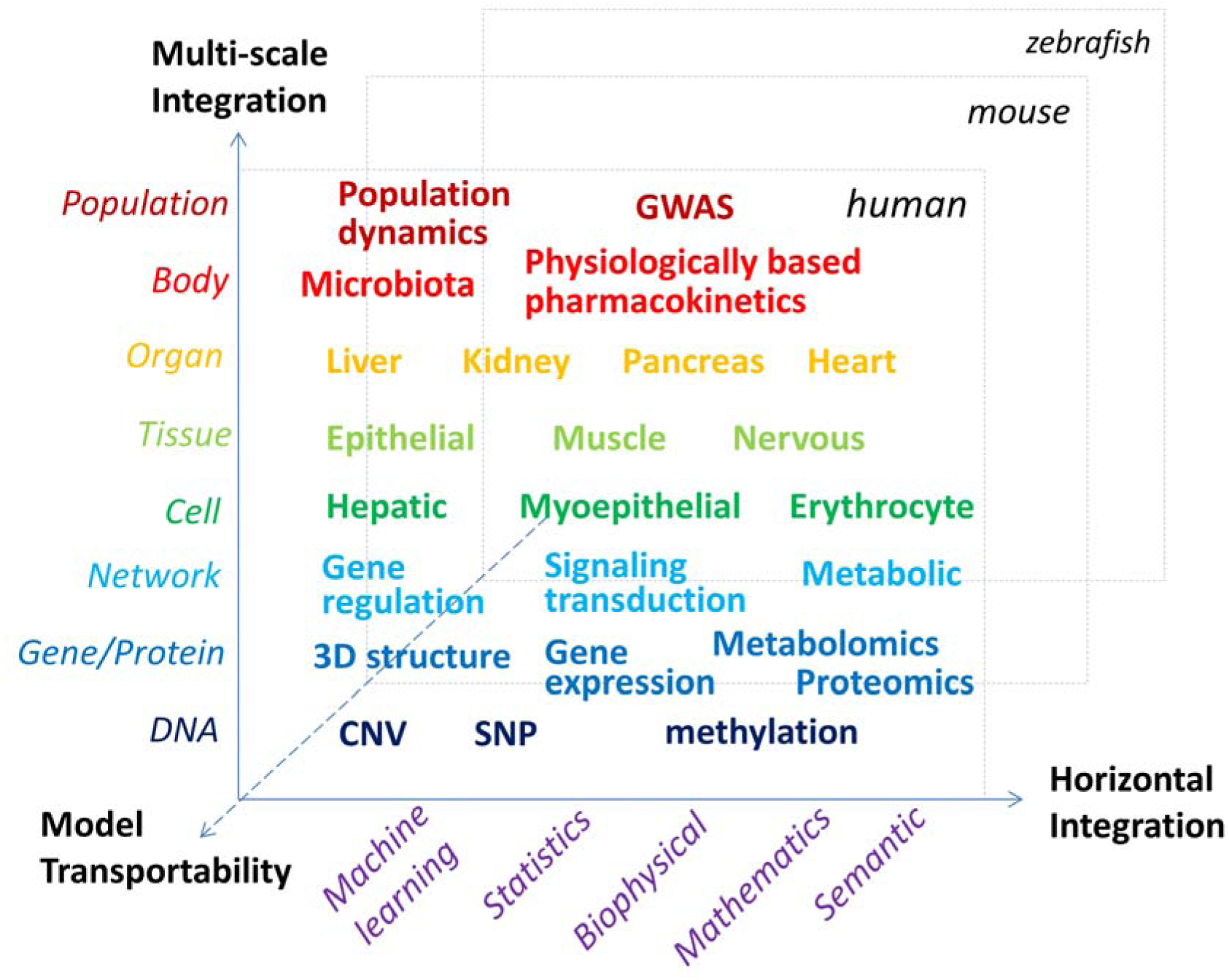
Model integration in systems pharmacology. Diverse models need to be integrated
across multiple methodologies, multiple heterogeneous data sets, organismal hierarchy, and species (transportability).

### Overview of existing techniques in big data analytics for horizontal model integration

Horizontal model integration can combine models that are built from different combinations of data sets, samples, features, and machine learning methods. Depending on training data and the base learner used, horizontal model integration can be cast as problems of ensemble learning, or data fusion in data science. Ensemble learning such as Bayesian Optimal Classifier (BOC), Bayesian Model Combination (BMC), Boosting, and Bootstrap Aggregation (Bagging) combines multiple models to hopefully form a better model using the same base learner, usually using a single data set. Data fusion techniques use multiple data sets that are from different data modalities (a.k.a. views) or sampled from features of a single modality. Data fusion methods can be roughly classified into three categories. In the first category, a model is built for each modality independently. Then these models are combined algorithmically, e.g., using stacked regression (54) or via a meta-predictor that has been widely used in chemoinformatics and bioinformatics, e.g., for predicting disease-associated mutations (55). The second category is to learn a joint representations of multiple data sets using deep neural networks (DNN) (56). The third category is to combine different data sets based on their semantic relationships. This can be achieved by four methods: multi-view learning (e.g., multi-kernel learning), similarity based method such as coupled collaborative filtering and manifold alignment, probabilistic dependency method such as Bayesian networks, and transfer learning (e.g., multi-target learning). These methods have been applied to address many problems in systems pharmacology modeling. For example, a predictive model based on multi-kernel learning is one of the best performers in the DREAM anti-cancer drug sensitivity challenge (57). Kernel-based matrix factorization that combines drug similarity and target similarity is a promising method to predict novel drug-target interactions (58). PARADIGM uses Bayesian network models to combine genetic variation, gene expression, and pathway information for gene enrichment analysis (59). Multi-task learning is proven to be a valuable approach for inferring multi-target QSAR models for lead optimization (60). The advances in big data technology provide new opportunities for model combination. As different models in the combination can be trained independently, parallel and distributed computing models such as MapReduce will facilitate the implementation of horizontal model integration (61; 62). In spite of these advances, new strategies are needed to address inherent problems in systems pharmacology modeling.

### Challenges in application of big data analytics to systems pharmacology

When adapting existing techniques in big data analytics to model integration in systems pharmacology, several challenges remain due to the inherent complexity of biological and clinical data. In addition to big volumes, pharmacological and clinical data are high-dimensional, incomplete and biased, heterogeneous, dynamic, and noisy. Firstly, unlike big data in other domains such as social networks, imaging processing, and natural language, where there are huge numbers of samples, pharmacological and clinical data may be sparse but high-dimensional. For example, in Genome-Wide Association Studies, the sequencing data of a whole genome can have hundreds of gigabytes with hundreds of thousands of Single Nucleotide Polymorphisms (SNPs). The sample size is typically in the range of hundreds of individual genomes. Although the total volume of data can be hundreds of terabytes, the number of variables is far larger than the number of samples. The One Million Genome Project cannot completely solve the under sampling problem due to the diversity of disease phenotypes and pharmacogenomics profiles. Handling extremely high-dimensional data is an unsolved problem in big data analytics. For example, without integrating with other data, GWAS data alone cannot identify disease-associated mutations (64). Secondly, existing data from pharmacology are often incomplete and biased. For example, only several thousand genes from multiple organisms have associated ligand binding information. Moreover, the number of associated ligands for each target is highly uneven, with many uncharacterized proteins playing important roles in drug action. Thirdly, in terms of heterogeneity, these data are across the hierarchical organization of an organism (molecule, pathway, cell, tissue, organ, patient, and population), across a wide spectrum of time scales, and across multiple species. As mentioned previously, multi-scale modeling is required (6). Even in the same organization and time scale of the same species, the data can be highly heterogeneous. For example, inter-tumor and intra-tumor heterogeneity have been observed ubiquitously (65). It is difficult to build a generalized machine learning model to predict anti-cancer drug sensitivity, as a new case can be unique and be out of the hypothesis space of trained models. Thirdly, the biological response to drug perturbation is dynamic. For example, cancer cells, bacteria and viruses can evolve rapidly to gain drug resistance. Systems pharmacology modeling should take the dynamics of drug response into account. Finally, in terms of noise, systems pharmacology must not only consider the signal to noise ratio of the various experimental methods and datasets, but also incorporate noise and stochasticity into its models, which is an intrinsic property of biological processes (66).

Conventional techniques for model integration in big data analytics are not sufficient to address the aforementioned challenges in systems pharmacology. In the case of ensemble learning, two of the most influential and practical methods, boosting and bagging, both have limitations. Boosting has been criticized by its incapability of dealing with noisy data sets (67). Original bagging algorithms based on random sampling of data further reduce the sample size for training, making the model building more difficult for high-dimensional data. Moreover, the heterogeneity of samples may represent the underlying functional space of biological system (e.g., different tissue types, pathogenicity, races etc.). The random sampling of data may not be the best strategy for bagging to handle heterogeneous biological or clinical data. Several recently developed techniques may offer new solutions to adapting the ensemble learning to model integration in systems pharmacology. For example, it is proven that the minimization of sample intersections in the bagged predictors will improve the performance of bagging (68). In other words, “bagging by design” that maximizes the model variance may outperform bagging at random. It implies that bagging may take advantage of the heterogeneity of data. Multi-view ensemble learning is another technique that may facilitate systems pharmacology modeling using high-dimensional data (69). Here, bagging, clustering, or random feature set partitioning etc. is applied to select multiple subsets of features. Each subset of features is used as a view to train a model. An ensemble is constructed by the combination of these models. As discussed elsewhere (10), the incorporation of biological knowledge into data-driven modeling is critical to the process of systems pharmacology modeling. For example, protein-protein interaction networks may assist the optimal feature set partitioning.

A fundamental challenge in big data analytics is to discover unknown-unknowns outside the existing domain of knowledge. It is particularly difficult for systems pharmacology. As mentioned previously, pharmacological and clinical data are often incomplete, biased, and heterogeneous. As a result, models built on these data only cover a portion of pharmacological and physiological space; that is, each model may be biased towards a certain knowledge domain. For example, druggable genes and proteins that have experimentally determined structures only partially overlap. Models built on these divergent data sets are complementary, but different. Existing paradigms for model integration may be not suitable. The underlying hypothesis of a voting strategy is that each model is better than random but not a strong performer. BMC finds the set of optimal weights that are based on observed data by randomly sampling the weight space. After training, the weight is fixed. It assumes that the high-weighted model always performs better than the low-weighted model. Both of these hypotheses do not hold in many cases of systems pharmacology, where a model could be strong for a case but weak to another one. If the case is outside the knowledge space covered by all models, it is possible that all models fail. Case-based reasoning (CBR) may provide an alternative solution to integrate heterogeneous models in systems pharmacology. In the field of artificial intelligence, CBR is an approach that solves a new problem by adapting solutions to a previously similar problem (70). When applying CBR to model integration, it may work as follows. Firstly, old cases are clustered based on the similarity among them. Secondly, the performance of each model is evaluated for each case cluster. This generates a performance matrix. Then, given a new case, its similarity to each cluster is calculated. Finally, the models are weighted using the similarity of their associated clusters to the query case and the performance matrix. It is noted that the model weight is case dependent in the CBR. The CBR strategy has been successfully applied to combine multiple protein-ligand interaction models for docking scoring, and significantly improve the performance of high-throughput screening (71). The challenge for CBR is how to select relevant features and how to assess the similarity between cases. The combination of advances in data science and domain-specific knowledge may provide feasible solutions (10).

The ultimate goal of systems pharmacology modeling is to generate novel hypothesis for discovering new knowledge and supporting decision making in drug discovery and clinical practice. Followed by experimental validation, new biological knowledge can be discovered by validating or refuting the hypothesis. In turn, this will increase the coverage of knowledge space, thereby facilitating systems pharmacology modeling. It is more demanding to apply systems pharmacology modeling to support decision making in drug discovery and clinical practice such as the prediction of anti-cancer drug efficacy for a particular patient. In such risk-sensitive domains, a reliable assessment of predictive modeling quality on an individual basis is essential. A fundamental assumption in machine learning is that the data sample on which an algorithm learns is representative of the complete data set to which the algorithm is applied. As a result, methods proposed to address prediction reliability include reverse transduction and local sensitivity analysis, bagging variance, local cross-validation, and local error modeling (72; 73). All, however, are tailored to generalize, and may not apply to an individual case that may fall outside the space of the training data. To assess the prediction reliability for a new case, it is critical to define the boundary of model space, and to determine if the new case falls within the model space. The CBR paradigm may provide a solution to this problem. Another strategy is to incorporate the systems pharmacology modeling into existing biological and clinical knowledge, as discussed in the following section.

## 4. Model interpretation and knowledge integration

The notable aim of systems pharmacology modeling is not only to maximize prediction accuracy but also to reveal the mechanism of drug action and to support decision making in drug discovery and clinical practice. Thus, it is necessary to have interpretable models, to integrate the model’s with existing knowledge, and to enable automated reasoning.

Systems pharmacology models can be assembled by either data-driven or mechanism-based approaches. These two approaches are often combined in systems pharmacology e.g., constraint-based modeling. Data-drive modeling, especially, machine learning approaches, often generate a black box. Recent works may facilitate opening the black box of machine learning models. For example, Random Prism has been proposed to be an alternative to widely used Random Forest methods (74). The base learner of Random Prism is the Prism algorithm that learns a set of IF-THEN rules instead of trees. It may provide a better representation of knowledge, which cannot be easily encoded as a decision tree. In another case, sequence motifs are extracted from the output of kernel-based learning algorithms (75).

Although mechanistic-based models offer a more straightforward explanation of drug action, challenges still remain to translate fragmented biophysical or mathematical descriptions into unified biological knowledge. Firstly, computational models should be coupled with existing biological and clinical knowledge. The coupling will allow us to evaluate the model, generate new hypotheses, or identify knowledge gaps. Secondly, a mechanistic understanding of drug actions requires the combination of molecular, network, phenotype, and other models. However, these models are developed independently at different scales and using different modeling languages. Thus it is not straightforward to integrate them into a unified model. Finally, mathematical languages used for modeling may not be easily comprehended by biologists and clinicians trying to establish causal relationships between genetic mutations, molecular interactions, as well as network modulations and pathophysiological processes and clinical outcomes. It is necessary to translate mathematical language into not only accessible human knowledge, but also a machine-readable representation for automated reasoning. To address these challenges, effective knowledge representations of input-output relationships from systems pharmacology modeling, which is both human comprehensive and machine readable, may be needed in addition to the development of ontologies that enable efficient communications between models as discussed in the previous section.

### Using semantic web for knowledge integration

The semantic web has promised to turn big data into linked and smart data, and has emerged as a powerful technique for knowledge integration in systems biology (76) and healthcare (77). In the semantic web, knowledge is represented by the Resource Description Framework (RDF) (http://www.w3.org/RDF/) as the form of Subject-Predicate-Object triplets. Domain knowledge is modeled as a graph of triplets. The graph model is stored in triple/RDF database management systems. The query language SPARQL has been developed to retrieval information from the triple/RDF store. The Web Ontology Language (OWL) has been proposed to support database queries and rule-based technologies.

With the maturing of semantic web technologies, they have been exploited to link heterogeneous datasets into a unified knowledgebase in systems biology. For example, BioGateway uses the semantic web to integrate the OBO foundry ontology, Gene Ontology, NCBI taxonomy, and UniProt (78). eXFrame provides a reusable framework for creating semantic web repositories of genomics experiments (79). Bio2RDF release 2 links 19 datasets (80). Many of them are directly relevant to systems pharmacology modeling, such as the Comparative Toxicogenomics Database (CTD) (81), DrugBank (82), Medical Subject Headings (MESH), National Drug Code Directory (NDC), Online Mednelian Inheritance in Man (OMIM) (83), Pharmacogenomics Knowledge Base (PharmGKB) (84), etc. It includes 57,850,248 unique subjects, 298,470,583 unique objects, 1,003 unique predicates, and 1,01,758,291 triplets. In addition, the semantic web has been applied to manage electronic health records (EHRs) with the aim to capture, standardize, integrate, describe, and disseminate health related information (85–88). It has been proposed that a semantic data-driven environment is needed to address big data challenges in health care (77). The consolidation of semantic systems biology and semantic health care may provide new opportunities for genome-wide association studies, pharmacogenomics, and personalized medicine.

### Integration of systems pharmacology modeling with biological and clinical knowledge

The efforts in applying the semantic web to systems biology and health care provide a solid foundation to advance the knowledge integration of systems pharmacology modeling. The incorporation of systems pharmacology modeling into a semantic rich knowledge base may harness the power of systems pharmacology modeling in generating novel hypotheses and support decision making. It is important to transform the quantitative results from predictive models into logical descriptions or rules between biological entities. Then the logical relationships can be represented as RDF triples. The subject and the object are biological concepts (entities) such as genes. The predicate is the molecular interaction, functional association, or causal relationships between biological entities. For example, a predicted interaction between a chemical A and an agonist conformation of receptor B can be represented as “A activate B” (Figure 3). An ontology-driven uniformed concept mapping is needed to link genes, proteins, biological pathways, and phenotypes as well as their interactions from diverse models. With the uniformed concept mapping based on the ontologies, and the translation of model outputs as an RDF triple, systems pharmacology models can be incorporated into existing semantic based systems biology and health care knowledge base. In this way, not only can software handle information to build inferences and test hypotheses, but also computational and mathematical models are human interpretable. An example is shown in Figure 3. A knowledge graph has the following triplets: Agonist and antagonist binding of PPAR up-regulate and down-regulate RAAS, respectively. The up-regulation of RAAS increases blood pressure, whereas its down-regulation decreases blood pressure. A computational model predicts that a drug binds the agonist or antagonist conformation of PPAR. The model result can be transfer to a triplet in the knowledgebase. Then the drug response phenotype (in this case, hypertension or hypotension) can be inferred.

**Figure 3.**
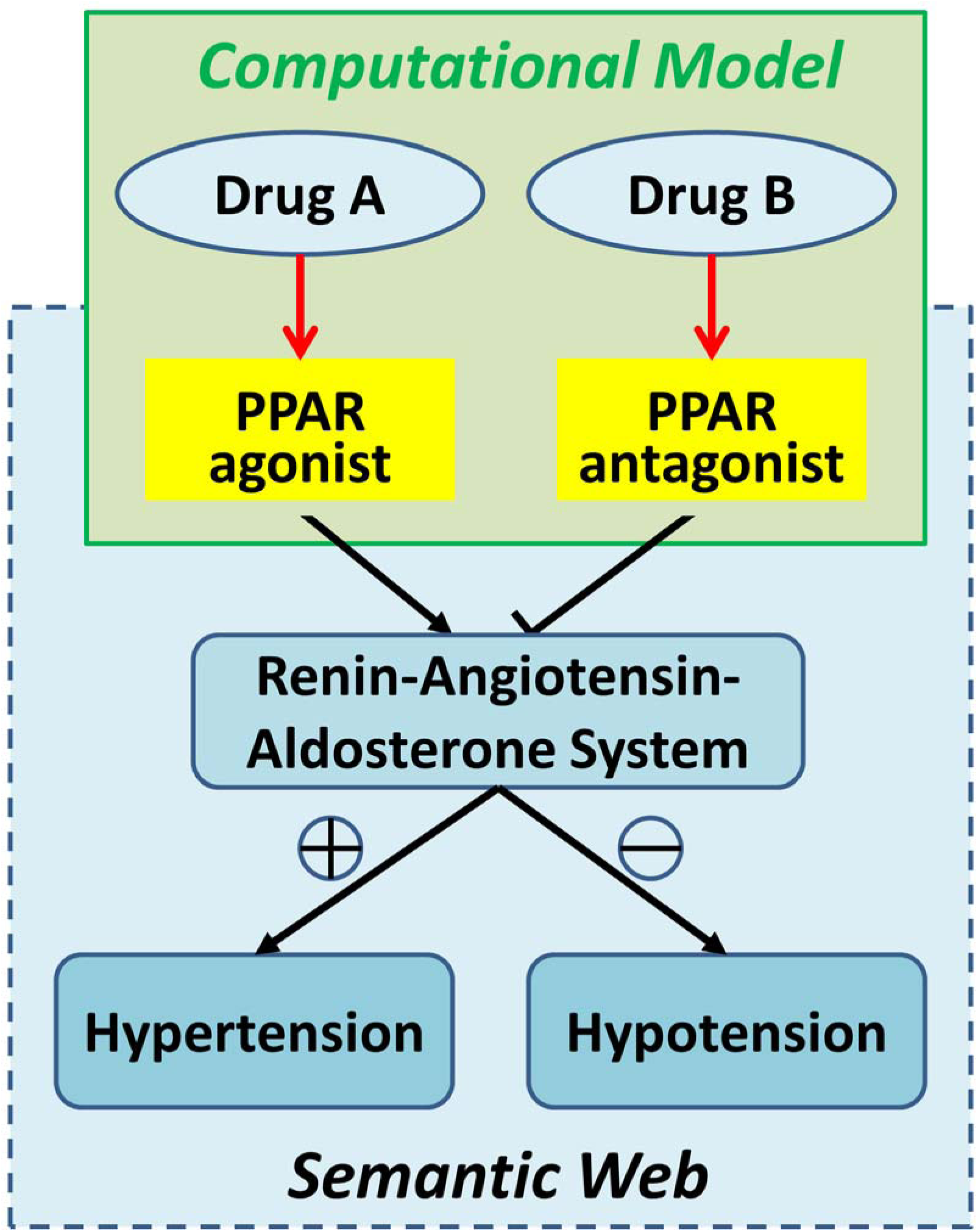
An example to integrate systems pharmacology modeling and the semantic web. The output of systems pharmacology models are translated into an RDF triplet and associated with a knowledge base that is built on semantic web technology. The knowledge base will support automated model validation, reasoning, and decision making.

## Conclusion

The conventional one-drug-one-target-one-disease drug discovery process has been less successful in tracking multi-gene, multi-faceted complex diseases. Systems pharmacology has emerged as a new discipline to tackle the current challenges in drug discovery. Systems pharmacology modeling utilizes diverse methodologies, integrates multiple omics data, crosses the hierarchy of an organism, spans a wide range of time scales, and addresses the uniqueness of the individual. Successful systems pharmacology modeling requires integrating multiple models to gain a holistic and comprehensive understanding of drug actions under diverse genetic and environmental conditions. Although data integration has already attracted tremendous attention in systems pharmacology, we face new challenges to enable systems pharmacology modeling to be findable, accessible, interoperable, reusable, reliable, interpretable, and actionable. Advances in big data technologies and data science may provide technical solutions to address these challenges. Beyond this, we need new business models to prompt open science that is essential for systems pharmacology. The NIH commons is an example to this direction.

## Acknowledgement

This research was supported by the National Library of Medicine of the National Institute of Health under the award number R01LM011986 (LX)

